# LPS-Induced Inflammation Reduces GABAergic Interneuron markers and Brain-derived Neurotrophic Factor in Mouse Prefrontal Cortex and Hippocampus

**DOI:** 10.1101/2023.11.15.567229

**Authors:** Sara Rezaei, Thomas D. Prévot, Erica Vieira, Etienne Sibille

## Abstract

Inflammation, reduced gamma-aminobutyric acidergic (GABAergic) function and altered neuroplasticity are co-occurring pathophysiologies in major depressive disorder (MDD). However, the link between these biological changes remains unclear. We hypothesized that inflammation induces deficits in GABAergic interneuron markers and that this effect is mediated by brain-derived neurotrophic factor (BDNF). We report here that systemic inflammation induced by intraperitoneal injection of lipopolysaccharide (LPS) (0.125, 0.25, 0.5, 1, 2mg/kg) in the first cohort of C57BL/6 mice (n=72; 9-12 weeks; 50% female) resulted in increased interleukin 1-beta and interleukin-6 in prefrontal cortex (PFC) and hippocampus (HPC), as measured using enzyme-linked immunosorbent assay (ELISA). Quantitative real-time polymerase reaction (qPCR) was used to explore the effect of LPS on the expression of GABAergic interneuron markers. In the PFC of the second cohort (n=39; 9-12 weeks; 50% female), 2mg/kg of LPS decreased the expression of somatostatin (*Sst*) (p=0.0014), parvalbumin (*Pv*) (p=0.0257), cortistatin (*Cort*) (p=0.0003), neuropeptide Y (*Npy*) (p=0.0033) and cholecystokinin (*Cck*) (p=0.0041), and did not affect corticotropin-releasing hormone (*Crh*) and vasoactive intestinal peptide (*Vip*) expression. In the HPC, 2mg/kg of LPS decreased the expression of *Sst* (P=0.0543), *C*ort (p=0.0011), *Npy* (p=0.0001), and *Cck* (p<0.0001), and did not affect *Crh, Pv,* and *Vip* expression. LPS decreased the expression of *Bdnf* in the PFC (P<0.0001) and HPC (P=0.0003), which significantly correlated with affected markers (*Sst, Pv, Cort, Cck,* and *Npy*). Collectively, these results suggest that inflammation may causally contribute to cortical cell microcircuit GABAergic deficits observed in MDD.

## 1. Introduction

Depression is a debilitating brain disorder affecting approximately 280 million people worldwide (Friedrich, 2017). Although the heterogeneity of depression has made it difficult to understand the underlying pathophysiology, depression is known to disrupt the gamma-aminobutyric acidergic (GABAergic) system, the main inhibitory neurotransmitter system (Fee et al., 2017). Clinical studies find reduced GABA levels in cerebrospinal fluid, plasma, occipital cortex, prefrontal cortex (PFC), and anterior cingulate cortex (ACC) of MDD subjects accompanied by reduced GABAergic neurotransmission as measured by transcranial magnetic stimulation-electromyography (Brambilla et al., 2003; Gerner et al., 1984; Gerner and Hare, 1981; Lefaucheur et al., 2008; Newton et al., 2019; Petty et al., 1990; Petty and Sherman, 1984; Radhu et al., 2013). While all GABAergic neurons share common elements, they differ in their molecular markers. The characterization of GABAergic neuron markers is now shedding light on GABAergic neuron-specific dysfunction in MDD.

The cortical microcircuit comprises diverse inhibitory GABAergic interneurons that regulate excitatory glutamatergic pyramidal neurons (PNs) and contribute to neuronal information processing (Fee et al., 2017; Tremblay et al., 2016)(Figure 1). GABAergic interneurons are classified by the expression of distinct molecular markers (neuropeptides or calcium-binding proteins), and the PN cellular compartment that they innervate (Tremblay et al., 2016). Three main subtypes of interneurons express the molecular markers somatostatin (SST), parvalbumin (PV), and vasoactive intestinal peptide (VIP) (Kubota, 2014). Corticotropin-releasing hormone (CRH), neuropeptide Y (NPY), cortistatin (CORT), and cholecystokinin (CCK) are used to further characterize interneuron subtypes (Tremblay et al., 2016). SST interneurons target the distal dendrites of PNs, and PV targets the perisomatic compartment of PNs. VIP interneurons mainly target SST interneurons resulting in the disinhibition of PNs at distal dendrites (Kubota, 2014). Reduced *SST* and genes co-expressed with *SST* (e.g. *CORT, NPY*) are reported in dorsolateral PFC, amygdala, and ACC of MDD subjects (Guilloux et al., 2012; Sibille et al., 2011; Tripp et al., 2011). PV expression is downregulated (Chung et al., 2018; Tripp et al., 2012) and VIP expression is mainly unchanged in MDD. The reduced GABAergic function in MDD results in decreased signal to noise ratio and dysfunction in information processing (Prévot and Sibille, 2021).

**Figure 1:**
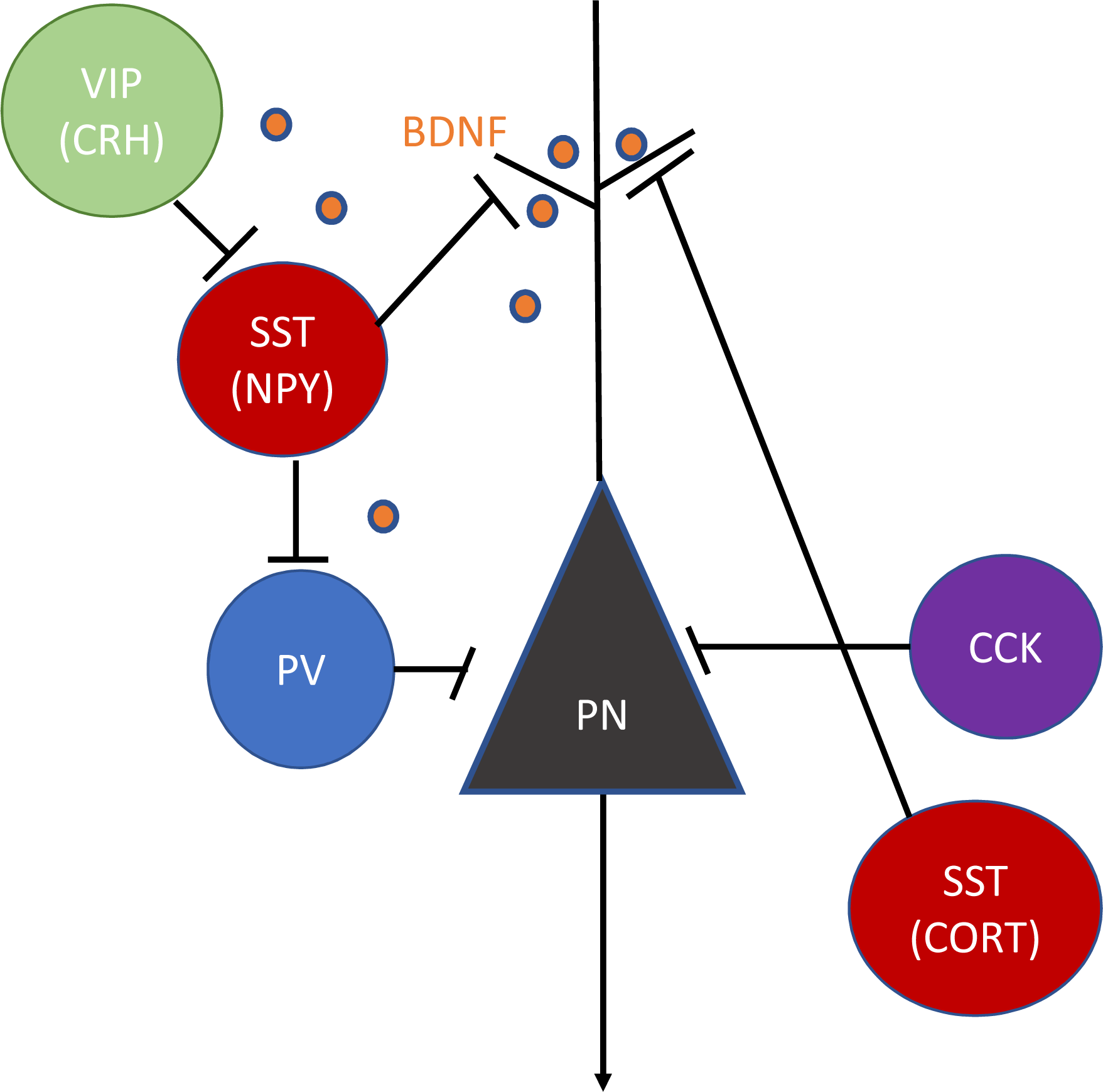
Cortical microcircuit schematic. Somatostatin (SST) interneurons provide tonic and phasic inhibition onto the pyramidal neuron (PN) dendrites, hence filtering incoming excitatory signals. Parvalbumin (PV) neurons inhibit the soma and axon initial segment of pyramidal neurons, the thalamocortical excitatory inputs by innervating. Upon stimulus, vasoactive intestinal peptide (VIP) neurons inhibit SST interneuron firing and induce disinhibition of PNs to allow information processing. Cholecystokinin (CCK) neurons mainly innervate the soma of PNs, providing strong rapid feedforward inhibition onto PNs. The neuropeptide corticotropin-releasing hormone (CRH) is mainly co-expressed with VIP interneurons and is important for providing inhibitory inputs to SST neurons. Neuropeptide Y (NPY) and cortistatin (CORT) are commonly co-expressed with SST interneurons. Brain-derived neurotrophic factor (BDNF) is a signaling neuropeptide mainly secreted from PNs and is important for dendritic compartments and GABAergic interneurons (adapted from Fee et al., 2017). Lines with black arrows indicate excitatory input. Lines with black endlines indicate inhibitory inputs.

In parallel to reduced markers of GABAergic neurons, postmortem studies show reduced pyramidal cell dendrites and synapses in PFC and hippocampus (HPC) (Kang et al., 2012; Morrison and Baxter, 2012; Rajkowska, 2000; Stockmeier et al., 2004), and reduced markers of neuroplasticity, namely reduced brain-derived neurotrophic factor (BDNF) in MDD (Ding et al., 2015; Guilloux et al., 2012; Oh et al., 2019; Thompson Ray et al., 2011; Tripp et al., 2011; Tripp et al., 2012). Genetic studies in rodents show that reducing BDNF levels leads to downregulations of various GABAergic markers (Oh et al., 2019), suggesting a contribution of reduced BDNF to the GABAergic interneuron phenotype in MDD.

In addition to GABAergic and neurotrophic deficits, inflammation is repeatedly associated with the pathophysiology of MDD, and conditions with underlying inflammation, such as obesity and diabetes, are risk factors for developing MDD (Raison et al., 2010). Several meta-analyses identify increased serum pro-inflammatory proteins, including c-reactive protein, tumour necrosis factor (TNF), and interleukin (IL)-6 (IL-6), in MDD subjects compared to healthy controls (Dowlati et al., 2010; Howren et al., 2009). There is higher interleukin (IL)-1beta in the cerebrospinal fluid of MDD subjects (Levine et al., 1999) and higher IL-6 levels in suicide attempters with MDD compared to controls (Lindqvist et al., 2009). Transcriptomic studies frequently report changes in inflammation-related pathways in MDD (Cho et al., 2019; Jansen et al., 2016; Leday et al., 2018). For instance, recent studies in the sgACC of post-mortem MDD cohort suggest the presence of sustained immune activation, such as cytokine response pathways linked to gene changes in dendritic targeting interneurons expressing CRH, VIP and SST (Shukla et al., 2022) and increased immune response genes, including IL-1beta in CRH expressing interneurons (Oh et al., 2022). All these studies suggest that the immune system may affect the GABAergic system and contribute to MDD.

In this study, we assessed whether specific GABAergic interneuron subtypes are preferentially affected by inflammation. We used bacterial endotoxin lipopolysaccharide (LPS), a cell wall component of gram-negative bacteria (Beutler, 2000) that activates toll-like receptor 4 (TLR4) found on innate immune cells and tested a putative direct causal effect on GABAergic interneurons. First, we hypothesized a causal link between inflammation and deficits in GABAergic interneuron markers. To address this, we investigated the expression levels of several GABAergic interneuron markers, *Sst, Crh, Vip, Pv, Npy, Cort*, and *Cck*, in the mouse PFC and HPC following LPS exposure for 18 hrs. Second, we hypothesized that altered BDNF mediates the effects of LPS-induced inflammation on GABAergic interneurons. To begin testing this, we measured *Bdnf* levels and calculated correlations between BDNF and GABAergic interneuron markers.

## 2. Material and methods

### 2.1 Animals

Eight-week-old male and female C57BL/6J mice (Jackson Laboratories, Bar Harbor, ME) were group housed in individually ventilated cages (IVC) and maintained on a 12-h light/dark cycle with food and water *ad libitum.* Mice underwent 2-weeks of habituation in the animal facility followed by three days of handling (Marcotte et al., 2021) to reduce their stress towards the experimenter. All procedures and experiments followed the Canadian Council on Animal Care guidelines and were approved by the Centre for Addiction and Mental Health (CAMH) Animal Care Committee. The first cohort (n=72; 9-12 weeks; 50% female) was used to measure inflammation and study the dose-dependent effects of LPS. The second cohort (n=39; 9-12 weeks; 50% female) was used to confirm results and extend it to other GABAergic neuron markers using a single dose of LPS.

### 2.2 Drug administration and brain dissection

Ultra-pure lipopolysaccharide from E.Coli 0111:B4 strain (InvivoGen, San Diego, CA) supplied as 5x10^6^ EU was diluted with 1ml of endotoxin-free water and aliquoted with sterile PBS and stored at -20°C. Mice were weighed and received one single intraperitoneal injection of the appropriate doses (0.125mg/kg, 0.25mg/kg, 0.5mg/kg, 1mg/kg, 2mg/kg) at 3PM, based on previous studies showing the doses are safe and induce behavioral and molecular changes (Wickens et al., 2018). After 18hrs, the mice were sacrificed by cervical dislocation, and the brains removed for dissection. This time point was selected because studies show that at 18 hrs, animals are past sickness-like behaviours (peaks at 2-6hrs) and experience depressive-like behaviours (peaks at 24hrs) (Dantzer et al., 2008). RNA-sequencing study finds that 1 day after LPS injection is the peak of differentially expressed genes in the brains of mice (Diaz-Castro et al., 2021). The olfactory bulb was cut off, revealing the anterior commissure. The subsequent coronal section contained the anterior forceps of the corpus collosum (AFCC), with the darker area in the middle representing the mPFC (the prelimbic and infralimbic cortex). A diamond shape was cut to remove the mPFC and avoid taking tissue from AFCC. The cortex on one hemisphere was gently peeled and pulled down to dissect the dorsal and ventral hippocampus. Both PFC and HPC were flash-frozen on dry ice.

### 2.3 Quantitative real-time PCR

PFC and HPC RNA and protein were extracted following the instructions of the Allprep RNA/protein kit (cataloge no. 80404; Qiagen, Hilden, Germany). RNA was measured by nanodrop, and cDNA was synthesized by mixing 200ng of total RNA with VILO reaction mixtures according to SuperScript VILO cDNA synthesis kit (cataloge no.11754050; Thermo Fisher Scientific, Waltham, MA) in a 20μL reaction. Ssco Advanced universal SYBR Green supermix (cataloge no.1725275; Bio-Rad, Hercules, CA) was used with qPCR to amplify cDNA of *Sst, Pv, Vip, Bdnf, Cck, Npy,* and *Cort*, on a Mastercycler real-time PCR machine (Eppendorf, Hamburg, Germany). The primer sets (IDT; Coralville, Iowa) for the transcripts and three internal controls are described in table 1. The three internal controls were beta-actin, cyclophilin A, and glyceraldehyde 3 phosphate dehydrogenase. Each qPCR run included three samples and eight transcripts of interest, including internal controls in quadruplicate using 96 well white shell/clear well plates. The cycle threshold (Ct) values were used to calculate delta Ct (dCt) for each reference gene by subtracting the three internal controls and calculating the geomean of dCt. The geomean dCt was used to calculate the relative expression level of each gene using the formula: relative expression level=2^-dCt*1,000. Relative expression level was converted to a percentage by the relative expression level/average expression level of the control group.

**Table 1:**
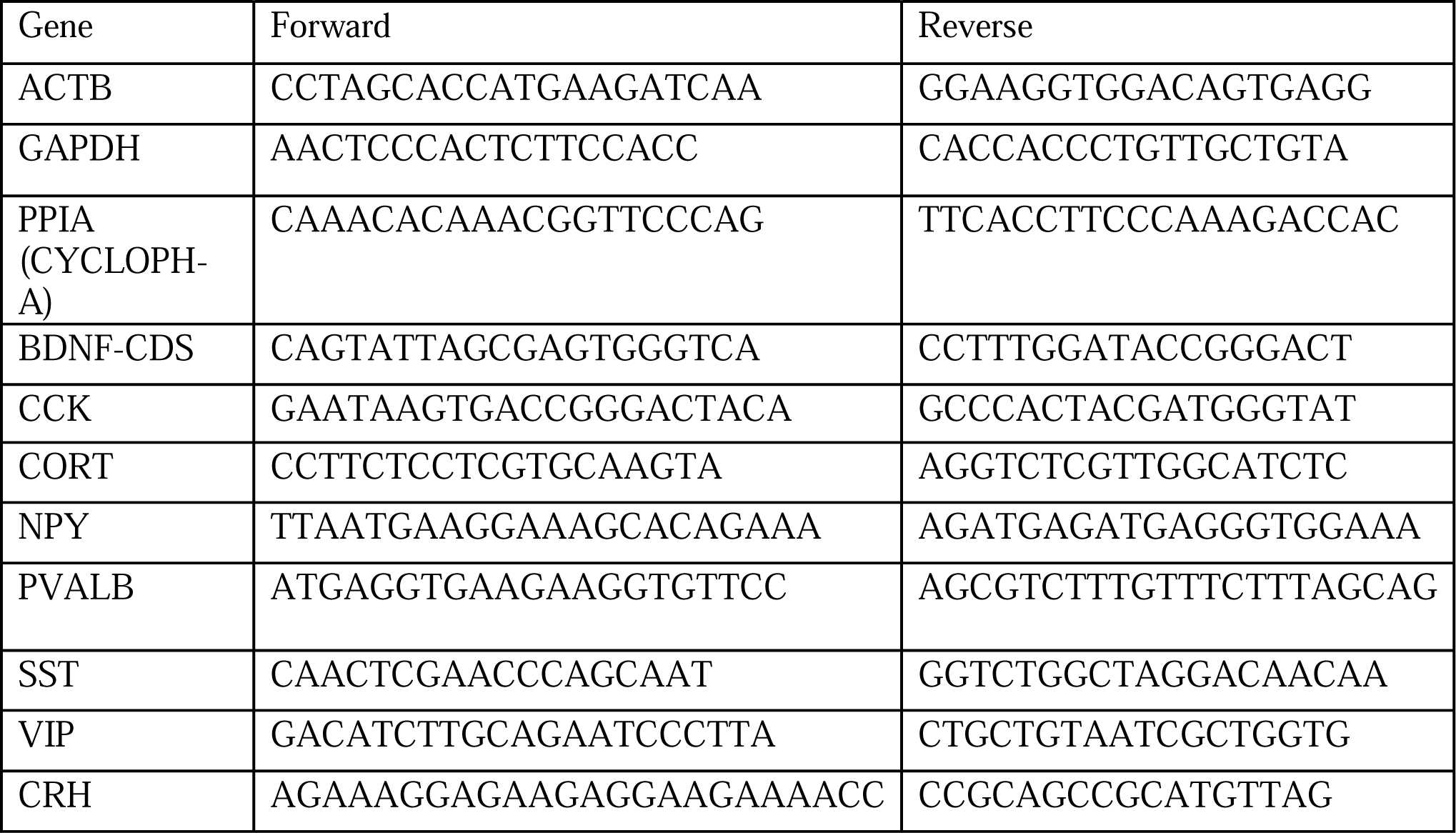
Primers used for qPCR.

### 2.4 Enzyme-linked immunosorbent assay (ELISA)

Bicinchoninic acid assay using PierceTM BCA Protein Assay Kit (cataloge no.23225, ThermoFisher, Waltham, MA) was used to measure the total extracted protein from PFC and HPC according to manufacturer’s instructions. To measure cytokines, DuoSet® ELISA kits for IL-6 (catalog no.DY406-05, R&D System, Minneapolis, Minnesota), TNF-alpha (catalog no.DY410-05, R&D System), and IL1-beta (catalog no.DY401-05, R&D System) were used in accordance with the manufacturer instructions. Briefly, a 96-well microplate was coated with 100μL solution containing diluted capture antibody in PBS for 18hrs. The plates were manually washed three times (350μL /well) with wash buffer (PBS-1X with 0.1%-Triton-X). The plate was blocked by 300μL of reagent diluent (1%-BSA in PBS) for 2hrs, followed by three washes. Cytokine standards and 1:2 diluted sample were added, followed by 18hrs of incubation at 4°C. The wash was repeated, and 100μL of detection antibody was added and incubated for 2hrs. After washing, 100μL of streptavidin-HRP was added to each well, and the plate was incubated for 30min at room temperature. The plate was washed, and 100μL of substrate solution (TMB) was added for 20min. Finally, 50μL of stop solution was added to each well, and the optical density was measured using a microplate reader set to 450nm (BioTek™ ELx 800™ Absorbance Reader, Agilent Technologies, Santa Clara, CA).

### 2.5 Statistical analysis

Statistical analysis was performed using GraphPad Prism 9.0. One-way or two-way ANOVA was used to determine the main effects of LPS, sex, and interaction. Appropriate post-hoc test, such as Dunnett’s, was performed. Mean group comparisons were done using Student’s T-tests, unpaired and two-tailed. Sexes were pooled when significant interactions with sex were not observed. Some samples were below the lower limit of quantification or were outliers, so were not included in the analyses. Pearson’s correlation was used to calculate correlations between expression of different genes.

## 3. Results

### 3.1 LPS dose-dependently increased pro-inflammatory cytokines IL1-beta and IL-6, with no effect on TNF-alpha in PFC and HPC

In response to LPS, an immune response is propagated into the brain and resident immune cells such as microglia synthesize inflammatory cytokines such as IL1-beta, IL-6, and TNF-alpha (Nonoguchi et al., 2022). To confirm the pro-inflammatory reaction in the LPS mice, protein levels of IL1-beta, IL-6, and TNF-alpha were measured in the PFC and HPC with increasing doses of LPS (0.125mg/kg, 0.25mg/kg, 0.5mg/kg, 1mg/kg, 2mg/kg) (Figure 2).

**Figure 2.**
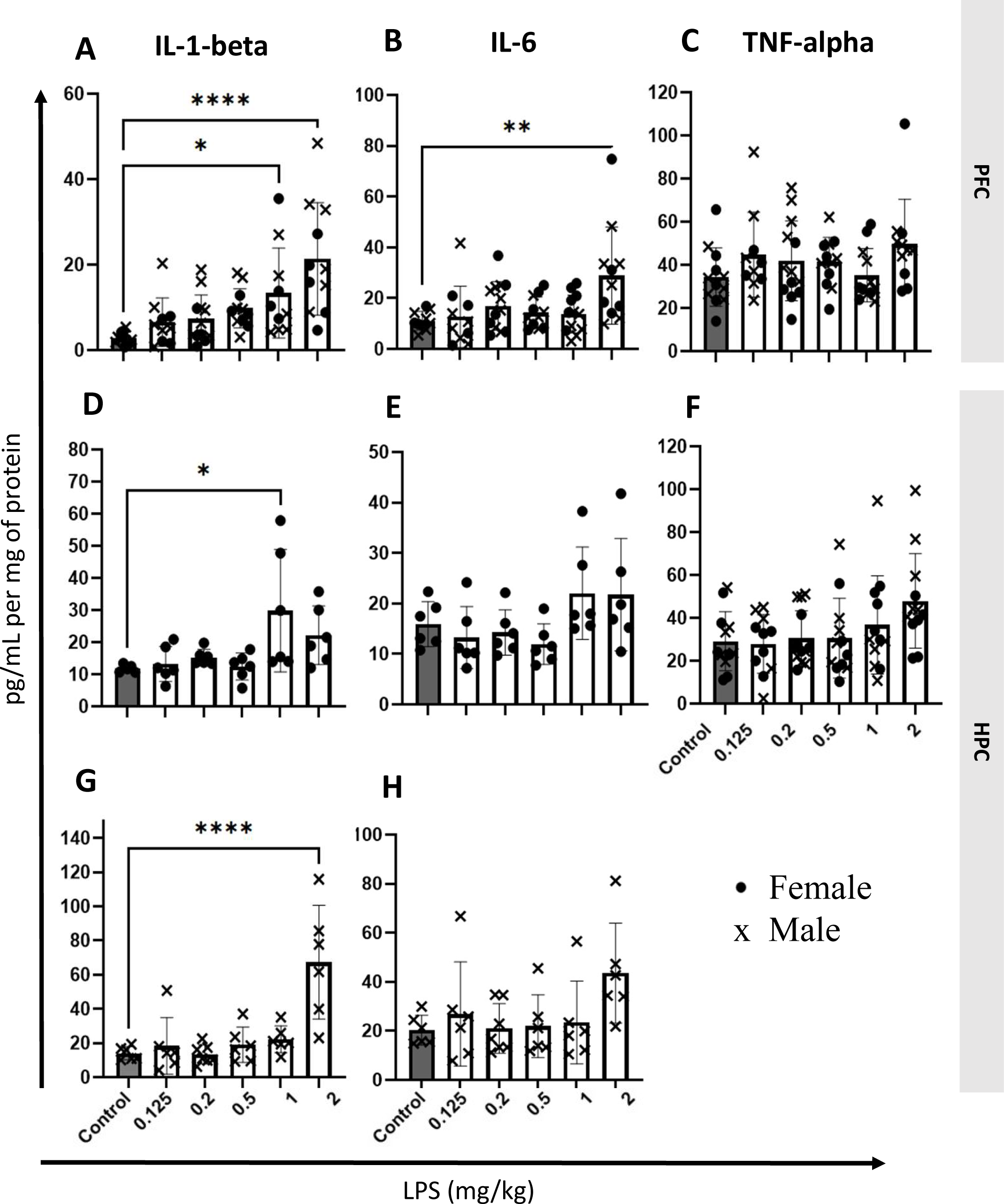
Effect of LPS (0.125mg/kg, 0.25mg/kg, 0.5mg/kg, 1mg/kg, 2mg/kg, i.p.) on protein levels of cytokines in prefrontal cortex (PFC) and hippocampus (HPC). **A** IL1-beta increased in PFC at 1mg/kg and 2mg/kg. **B** IL-6 increased in PFC at 2mg/kg. **C** TNF-alpha did not change in PFC. **D** & **G** IL1-beta increased in female HPC at 1mg/kg and in male at 2mg/kg. **E** & **H** IL-6 did not change in the HPC of females, nor males. **F** TNF-alpha did not change in HPC. Results are expressed as individual animals and mean ± SEM. Females are shown as circles and males as x symbol. *p<0.05, **p<0.01, ***p<0.001 and ****p<0.0001 as compared to control group.

#### 3.1.1 Prefrontal cortex

LPS significantly increased IL1-beta levels (F_5,53=_7.825; P<0.0001) with no effect on sex (F_1,53=_2.115; P<0.1517) or interaction (F_5,53=_1.397; P<0.2403). IL1-beta increased at LPS doses 1mg/kg (p=0.0135), and 2mg/kg (p<0.0001) compared to controls (Figure 2A). LPS significantly increased IL-6 levels (F_5,55=_3.690; P=0.0060) with no effect on sex (F_1,55=_0.5454; P=0.4634) or interaction (F_5,55=_0.4297; P=0.8260). IL-6 increased at LPS dose 2mg/kg (p=0.0019) compared to controls (Figure 2B). LPS did not significantly affect TNF-alpha levels (F_5,58=_1.423; P=0.2294) (Figure 2C).

#### 3.1.2 Hippocampus

LPS significantly increased IL1-beta levels (F_5,60=_10.23; P<0.0001) with significant effect on sex (F_1,60=_7.004; P<0.0104), and interaction (F_5,60=_6.063; P=0.0001). Main effect of LPS on IL1-beta were observed in female (p=0.0142, Figure 2D) and male (p<0.0001, Figure 2G). IL1-beta increased in female at 1mg/kg dose (p=0.0135) and in male at 2mg/kg (p<0.0001). LPS significantly increased IL-6 levels (F_5,58=_5.919; P=0.0002), with significant effect on sex (F_1,58=_8.468; P=0.0051), and interaction (F_5,58_=2.569; P=0.0362), although main effect of LPS on IL-6 levels were only at trend levels when split by sex (females, p=0.0725, Figure 2E; males, p=0.1067, Figure 2H). LPS did not significantly affect TNF-alpha levels (F_5,60=_2.252; P=0.0606) (Figure 2F).

### 3.2 LPS dose-dependently decreased *Sst* and *Pv* expression, with no changes on *Crh* and *Vip* in the PFC

LPS significantly decreased *Sst* expression (F_5,60_=2.580; P=0.0352), with no effect on sex (F_1,60_=1.388; P=0.2433) or interaction (F_5,60_=0.2412; P=0.9426). In two-group comparisons, Sst decreased at doses 0.125mg/kg (p=0.0316), 0.2mg/kg (p=0.0235), 0.5mg/kg (p=0.0084), and 1mg/kg (p=0.0366) (Figure 3A). LPS significantly decreased *Pv* expression (F_5,60_=4.133; P=0.0027), with no effect on sex (F_1,60_=1.441; P=0.2346) or interaction (F_5,60_=0.4855; P=0.7857) (Figure 3B). In two-group comparisons, *Pv* decreased at doses 0.2mg/kg (p=0.0006) and 1mg/kg (p=0.0018). LPS did not significantly affect *Crh* (F_5,59_=0.2775; p=0.9237, Figure 3C) or *Vip* expression (F_5,59_=1.354; P=0.2549, Figure 3D).

**Figure 3.**
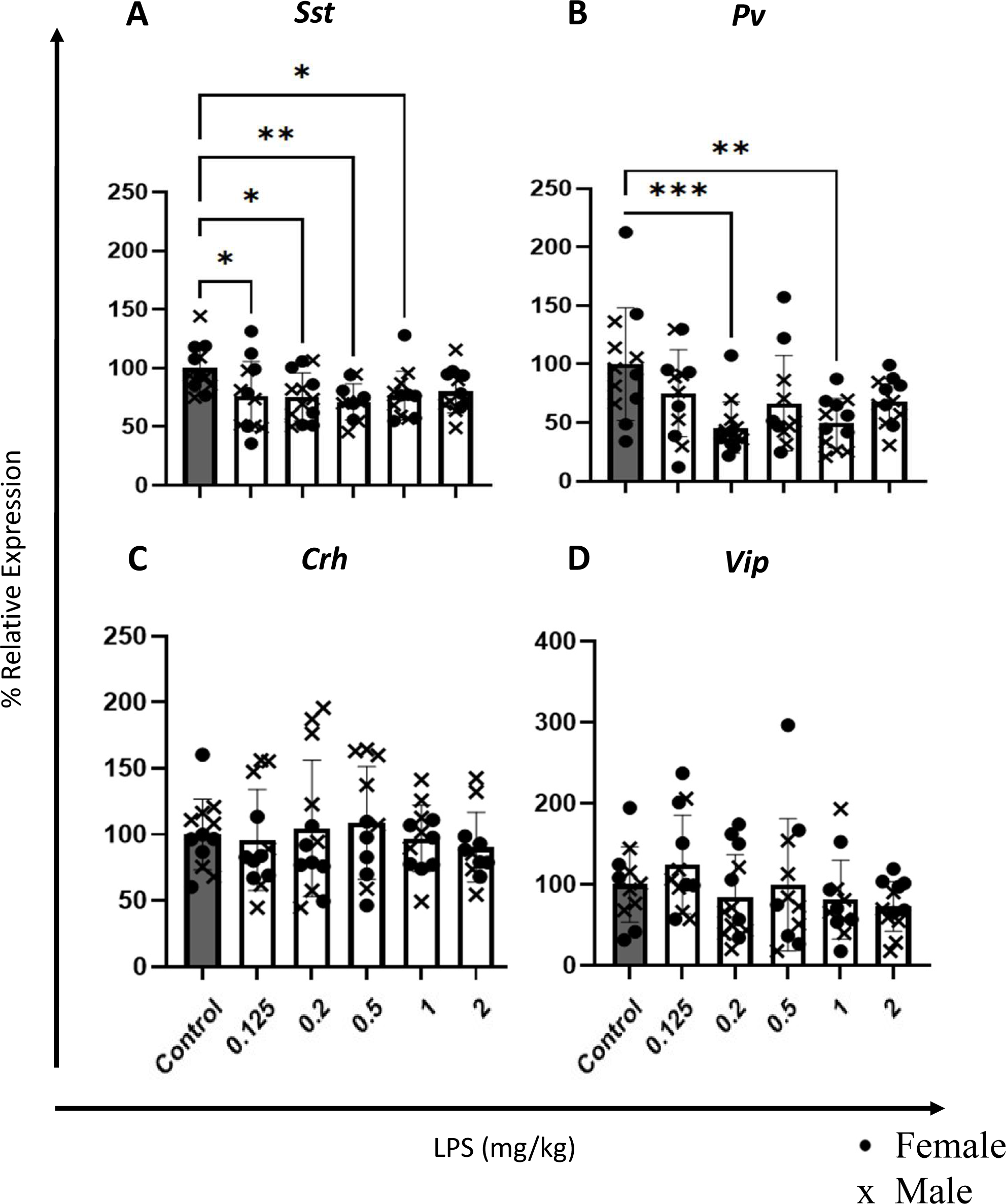
Effect of LPS (0.125mg/kg, 0.25mg/kg, 0.5mg/kg, 1mg/kg, 2mg/kg, i.p.) on the gene expression of *Sst*, *Pv, Crh,* and *Vip* in the prefrontal cortex (PFC). **A** *Sst* decreased at 0.125mg/kg, 0.25mg/kg, 0.5mg/kg, and 1mg/kg. **B** *Pv* decreased at 0.25mg/kg and 1mg/kg. **C** *Crh* did not change. **D** *Vip* did not change. Results are expressed as individual animals and mean ± SEM. Females are shown as circles and males as x symbol. *p<0.05, **p<0.01, ***p<0.001 and ****p<0.0001 as compared to control group.

### 3.3 LPS dose 2mg/kg decreased *Sst, Pv, Cort, Npy* and *Cck* expression with no changes on *Crh* and *Vip* in PFC

A larger independent cohort of male and female mice was next used to confirm the findings and investigate additional interneuron markers (Figure 4). The effect of a single 2mg/kg dose of LPS was selected for this cohort, based on the LPS-dose dependent cohort results. *Npy* and *Cort* expression were measured as additional markers co-expressed mainly with *Sst*. *Cck* expression was also measured, as a neuropeptide expressed in a different population of interneurons, which mainly target PNs directly. LPS significantly decreased *Sst* expression (F_1,35_=11.97; P=0.0014) with no effect on sex or interaction, and *post-hoc* analyses revealed 2mg/kg dose decreased *Sst* levels by 20% (p=0.0021) compared to controls (Figure 4A). LPS significantly decreased *Pv* expression (F_1,33_=5.455; P=0.0257) with no effect on sex or interaction, and *post-hoc* analyses revealed 2mg/kg dose decreased *Pv* levels by 20% (p=0.0290; Figure 4B). LPS did not affect *Crh* expression (F_1,35_=0.4637; P=0.5004; Figure 4C) or *Vip* expression (F_1,34_=0.6873; P=0.4129; Figure 4D). LPS significantly decreased *Cort* expression (F_1,34_=16.48; P=0.0003) with significant effect on sex (F_1,34_=4.218; P=0.0477), and interaction (F_1,34_=4.218; P=0.0477). In females, LPS did not affect *Cort* expression significantly (p=0.1011; Figure 4E) but decreased it in males by 43% (p=0.0012; Figure 2F). LPS significantly decreased *Npy* expression (F_1,34_=9.956; P=0.0033) with no effect on sex or interaction, and *post-hoc* analyses revealed 2mg/kg dose decreased *Npy* levels by 21% (p=0.0025; Figure 4G). LPS decreased *Cck* expression (F_1,34_=9.502; P=0.0041) with no effect on sex or interaction, and *post-hoc* analyses revealed 2mg/kg dose decreased *Cck* levels by 22% (p=0.0030; Figure 4H).

**Figure 4.**
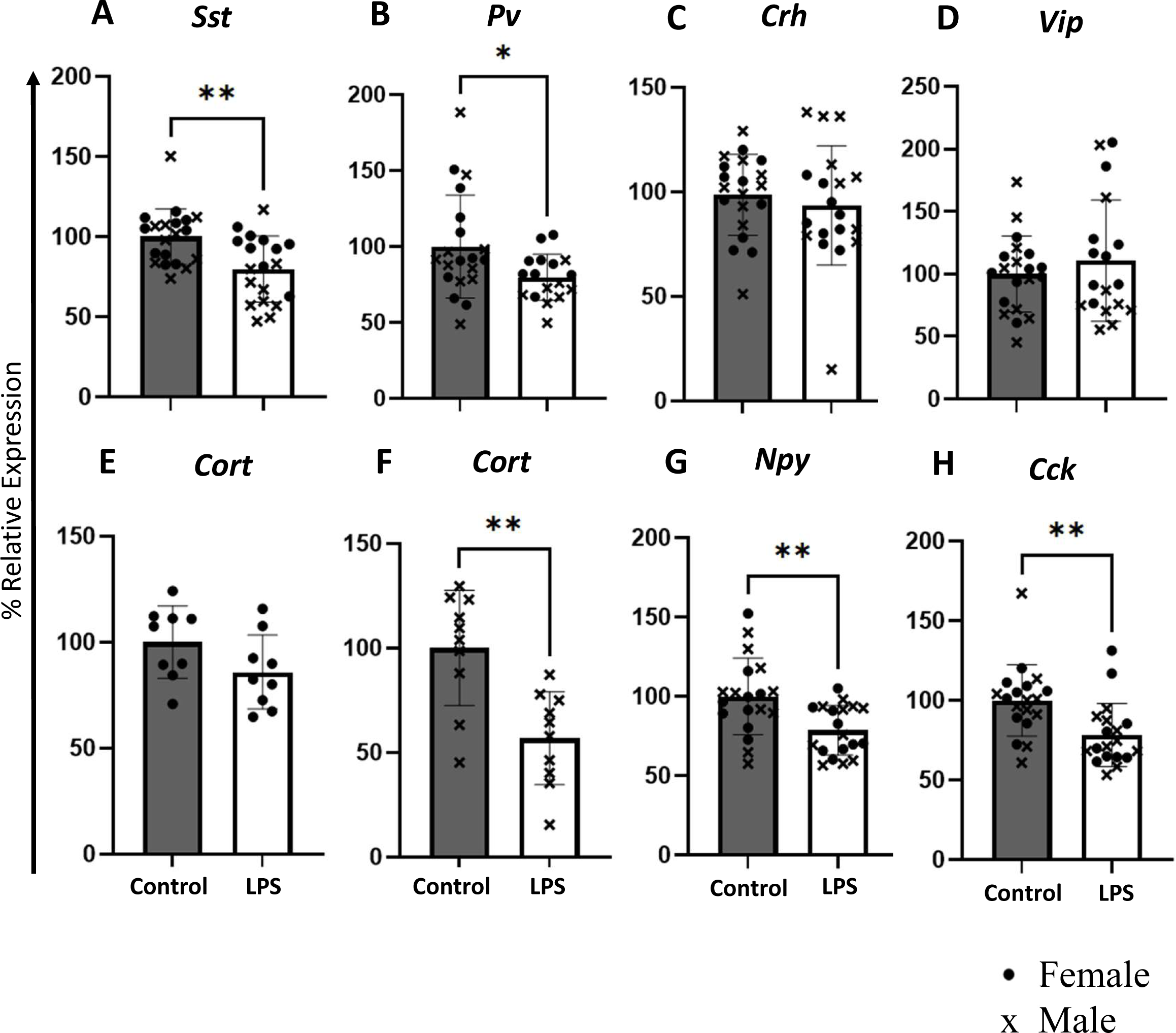
Effect of 2mg/kg of LPS on the expression of the *Sst, Pv, Crh*, *Vip, Cort, Npy,* and *Cck* genes in the prefrontal cortex (PFC). **A** *Sst* levels decreased. **B** *Pv* levels decreased. **C** *Crh* levels did not change. **D** *Vip* levels did not change. **E-F** *Cort* did not change in females and decreased in males. **G** *Npy* levels decreased. **H** *Cck* levels decreased. Results are expressed as individual animals and mean ± SEM. Females are shown as circles and males as x symbol. *p<0.05, **p<0.01, ***p<0.001 and ****p<0.0001 as compared to control group.

### 3.4 LPS dose 2mg/kg decreased *Sst, Cort, Npy* and *Cck* expression, with no changes on *Crh, Pv,* and *Vip* in the hippocampus

LPS decreased *Sst* expression (F_1,34_=3.974; P=0.0543) with no effect on sex or interaction, and *post-hoc* analyses revealed 2mg/kg dose decreased *Sst* levels by 12% (p=0.0616) compared to controls (Figure 5A). LPS had no effect on *Pv* (F_1,32_=0.8099; P=0.3749), *Crh* (F_1,34_=0.4760; p=0.4949), or *Vip* expression (F_1,32_=0.0356; P=0.8516) (Figures 5B-D). LPS decreased *Cort* expression (F_1,34_=12.70; P=0.0011) with no effect on sex or interaction, and *post-hoc* analyses revealed 2mg/kg dose decreased *Cort* levels by 23% (p=0.0008; Figure 5E). LPS decreased *Npy* expression (F_1,33_=18.34; P=0.0001), with no effect on sex or interaction, and *post-hoc* analyses revealed 2mg/kg dose decreased *Npy* levels by 28% (p=0.0002; Figure 5F). LPS decreased *Cck* expression (F_1,34_=29.46; P=0.0001) with no effect on sex or interaction, and *post-hoc* analyses revealed 2mg/kg dose decreased *Cck* levels by 35% (p=0.0001; Figure 5G).

**Figure 5.**
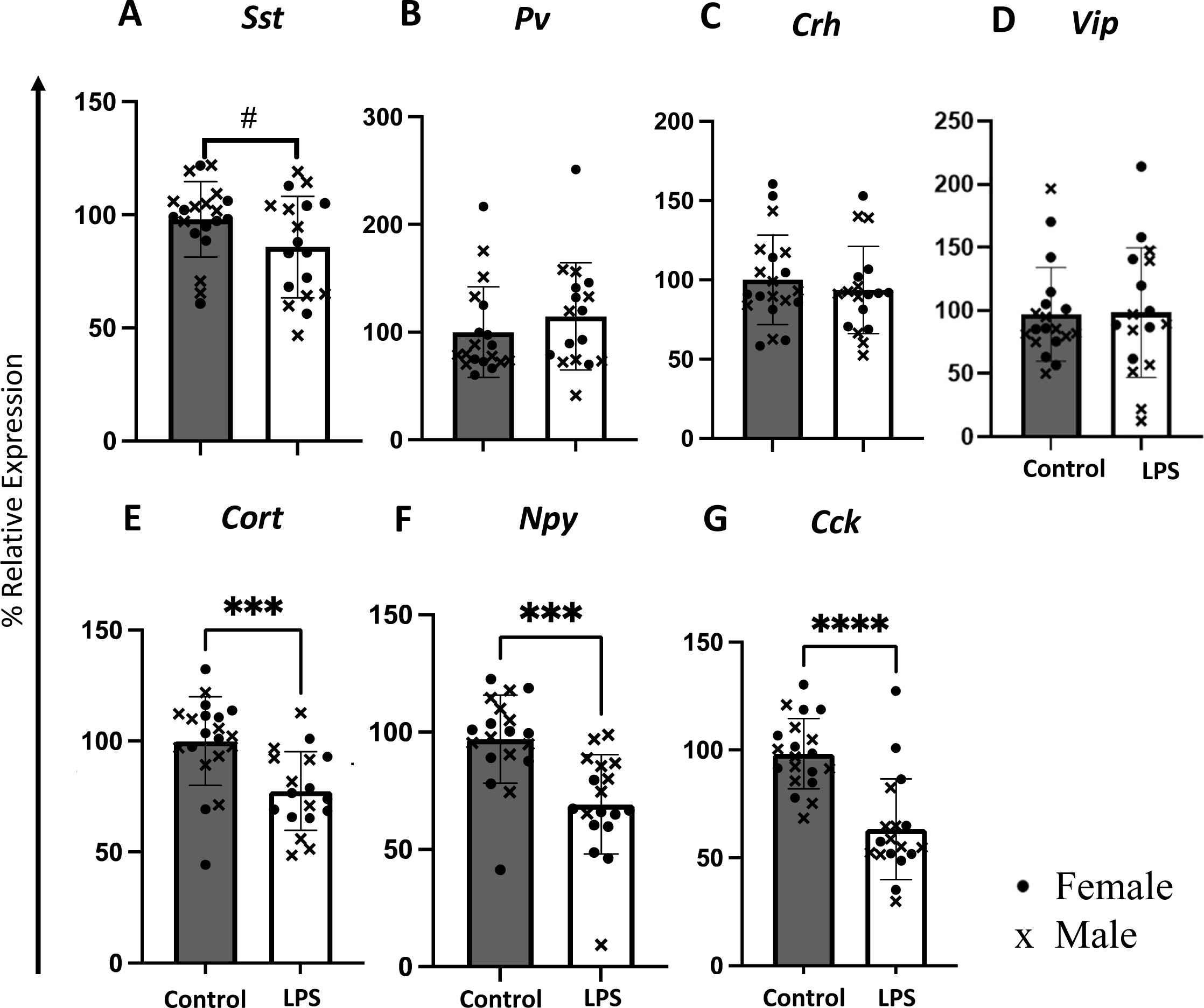
Effect of 2mg/kg of LPS on the expression of the *Sst, Pv, Crh, Vip, Cort, Npy,* and *Cck* genes in the hippocampus (HPC). **A** *Sst* levels had a trend towards decrease. **B-D** *Pv*, *Crh* and *Vip* levels did not change. **E** *Cort* levels decreased. **F** *Npy* levels decreased. **G** *Cck* levels decreased. Results are expressed as individual animals and mean ± SEM. Females are shown as circles and males as x symbol. *p<0.05, **p<0.01, ***p<0.001 and ****p<0.0001 as compared to control group.

### 3.5 Investigating a putative role for *Bdnf* in LPS-induced GABAergic interneuron marker changes

In the PFC, 2mg/kg of LPS significantly reduced *Bdnf* expression (F_1,35_=60.56; P<0.0001) with significant effect on sex (F_1,35_=5.612; P=0.0235), and quantitative interaction (F_1,35_=5.612; 0.0235), resulting in a 56% decrease in males (p<0.0001; Figure 6A) and 30% decrease in females (p=0.0046, Figure 6B). In the HPC, LPS decreased the expression of *Bdnf* (F_1,34_=16.58; P=0.0003), with no effect on sex (F_1,34_=0.3564; P<0.5545) or interaction (F_1,34_=0.3564; P<0.5544) and *post-hoc* analyses revealed 2mg/kg dose decreased *Bdnf* levels by 24% (p=0.0002) compared to controls (Figure 6C). Next, Pearson correlation (r) were assessed between gene expression of *Bdnf* and GABAergic interneuron markers (*Crh, Sst, Pv, Vip, Cck, Cort, Npy*). Significant positive correlations were observed between *Bdnf* and, in decreasing order of magnitude, *Cort* (r=0.48; p<0.0001), *Npy* (r=0.46; p<0.0001), *Sst* (r=0.41; p<0.0001), *Pv* (r=0.33; p<0.0001), and *Cck* (r=0.35; p=0.0018). No significant correlations were observed for *Crh* or *Vip* (Figure 6D).

**Figure 6.**
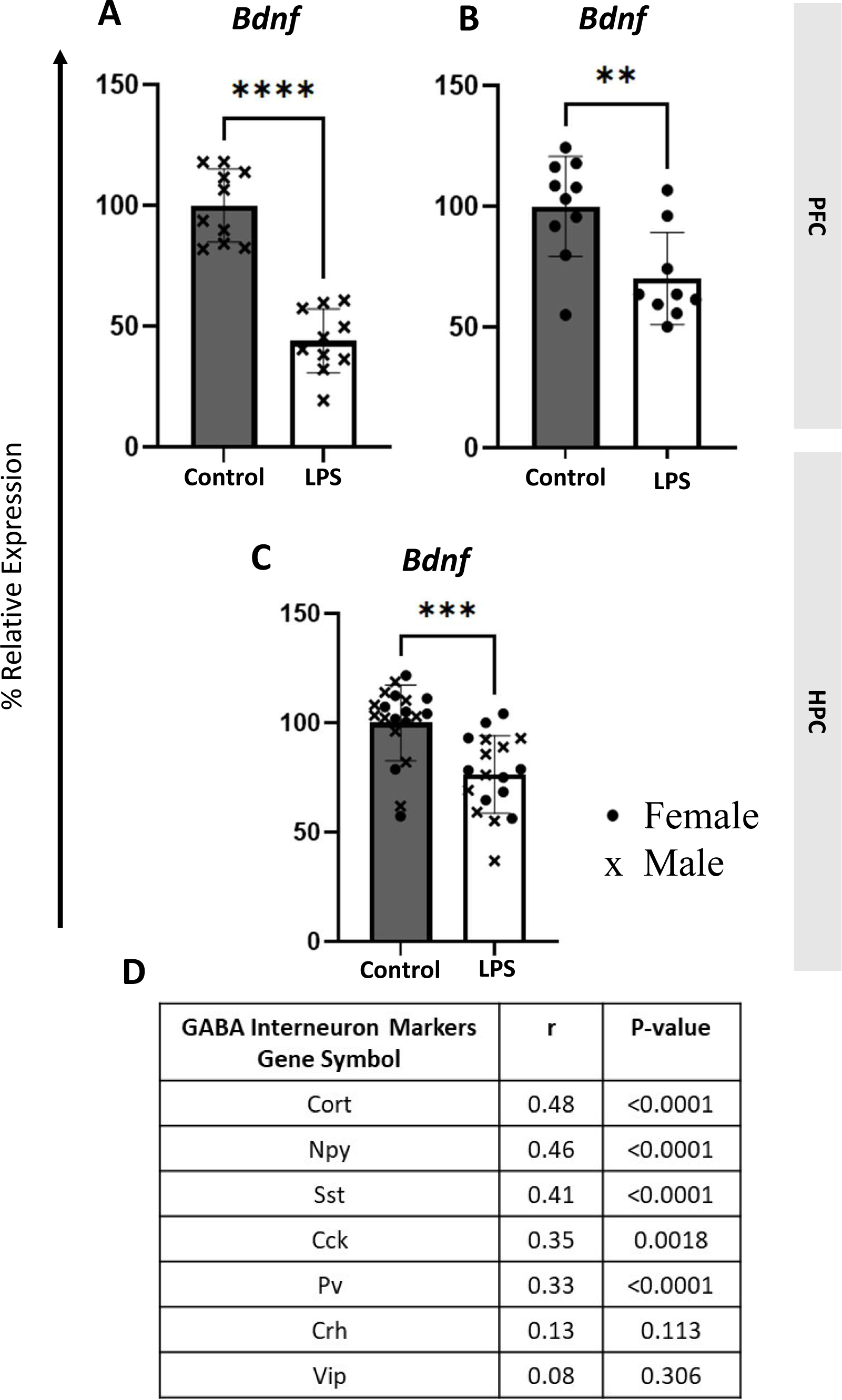
Effect of 2mg/kg of LPS on the expression of *Bdnf* gene in PFC and HPC and correlations with GABAergic interneuron marker genes. **A-B** 2mg/kg LPS decreased *Bdnf* levels in the PFC of males and females. **B** LPS 2mg/kg decreased *Bdnf* levels in the HPC. **E** *Bdnf* shows high correlation to *Cort*, *Npy,* and *Sst,* moderate correlations with *Cck* and *Pv,* and no correlation with *Crh* and *Vip.* Results are expressed as individual animals and mean ± SEM. Females are shown as circles and males as x symbol. *p<0.05, **p<0.01, ***p<0.001 and ****p<0.0001 as compared to control group.

## 4. Discussion

### 4.1 Selective vulnerability of interneurons to LPS-induced inflammation

Little is known about the role of inflammation on the disrupted functioning of GABAergic interneurons, a cellular phenotype that correlates with and potentially contributes to MDD. In this study, we investigated a putative causal link between inflammation and deficits in GABAergic interneurons, and report that LPS-induced inflammation reduced selective markers of GABAergic interneurons. *Bdnf,* a neurotrophic factor relevant to interneuron plasticity, was similarly downregulated by LPS and correlated with the expression of GABAergic markers affected by LPS.

Our results show that LPS-induced inflammation decreased the expression of three dendritic targeting interneuron neuropeptides *Sst, Cort*, and *Npy* which are indices of SST interneuron function, supporting a selective vulnerability of SST neurons to inflammation (Tomoda et al., 2022). While, the functional effects of *Sst* deficits on the neuron and at microcircuit level are beyond the scope of this study, *Sst* reduction or ablation studies in mice find similar molecular changes, including reduced *Bdnf,* GABA-related genes such as *Gad67, Cort* and increased emotionality (Lin and Sibille, 2015; Soumier and Sibille, 2014) that echo what is observed in human MDD. LPS-induced inflammation also affected *Pv* and *Cck*, as markers of interneurons that target the perisomatic region of PNs. Other studies with the LPS model in rats find reduced *Pv* and *Gad67* expressions in medial PFC (Ji et al., 2020) and reduced *Sst* expression in the hippocampus (Gavilán et al., 2007). We find that *Sst* and *Pv* are only significantly affected in the PFC (Figure 4), possibly due to the more robust inflammatory response, as evidenced by higher levels of pro-inflammatory cytokines IL1-beta and IL-6 in the PFC compared to the HPC (Figure 2). The blood-brain barrier of PFC is shown to be more vulnerable to disruption by the inflow of peripheral inflammatory mediators than other regions (Erickson and Banks, 2011; Nonoguchi et al., 2022). The more robust inflammatory response in the PFC may suggest that PFC interneurons are either under greater inflammatory insult or more vulnerable to LPS immune challenge that induces an immune response peripherally and centrally. Using blood sample from LPS-induced mice, future studies can correlate cytokine levels in the periphery with markers in the brain. Moreover, *Cort* and *Bdnf* expression displayed sex x LPS interaction effects in the PFC of mice, in conjunction with a stronger increase in IL-beta levels in male HPC compared to females. Similarly, a stronger neuroinflammatory response as measured by IL1-beta and TNF-alpha is found in male mice compared to females (Nonoguchi et al., 2022; Rossetti et al., 2019), suggesting that sex differences in the immune response are mediating the effects on *Cort* and *Bdnf*.

This study is the first to investigate the effect of LPS-induced inflammation on *Cck*, finding that its expression is significantly reduced in PFC and HPC. Cck protein decreases in the plasma and hypothalamus of LPS-treated mice (Weiland et al., 2005). There are limited studies on CCK neurons in MDD (Banasr et al., 2017). However, this study suggests that CCK interneurons are vulnerable to inflammation, and the effect of their deficits on the cortical microcircuit requires future investigation.

VIP interneurons are unaffected by LPS-induced inflammation as there was no change in the expression of *Vip* nor *Crh*, markers mainly co-expressed by VIP interneurons (Oh et al., 2022). Few studies have explored the extrahypothalamic CRH and its function in inhibiting PNs through GABA synthesis (Chen et al., 2020; Dedic et al., 2018; Kubota, 2014). CRH-expressing cells are reduced in ACC, dorsolateral PFC, and amygdala of MDD subjects, and it is suggested that the GABAergic function of CRH-expressing interneurons is impaired (Ding et al., 2015; Oh et al., 2022). Previous studies find that LPS increases *Crh* levels in the hypothalamus (Ribot et al., 2003). However, no changes in *Crh* expression in the PFC and HPC are found, suggesting differential vulnerability between cortical and subcortical *Crh* neurons to inflammation.

Our results show that LPS affects the cortical microcircuit and changes the same profile of GABAergic interneurons (*Sst, Npy, Cort, Pv*) that are reduced in MDD (Figure 7). GABAergic deficits and inflammation are also associated with other neuropsychiatric disorders. Human post-mortem studies find reduced *SST,* and *NPY* in schizophrenia (Guillozet-Bongaarts et al., 2014; Mellios et al., 2009; Volk et al., 2012), and bipolar disease (Konradi et al., 2004; Sibille et al., 2011). PV is also reduced in schizophrenia (Beasley et al., 2002; Mellios et al., 2009; Volk et al., 2012), and bipolar disease (Pantazopoulos et al., 2007). Like MDD, there is upregulated immune/inflammation-related genes in the hippocampus and cerebral cortex of patients with schizophrenia (Fillman et al., 2013; Hwang et al., 2013) and chronic-low grade inflammation is associated with bipolar disease (Barbosa et al., 2014). RNA-sequencing data from the HPC of patients with schizophrenia show that immune/inflammation response genes are associated with deficits in GABAergic interneurons, particularly with the PV neurons (Kim et al., 2016). Thus, our findings are applicable to neuropsychiatric disorders with GABAergic system deficits and suggest that an underlying chronic inflammation may directly affect selective GABAergic interneurons to reduce their marker expression.

**Figure 7.**
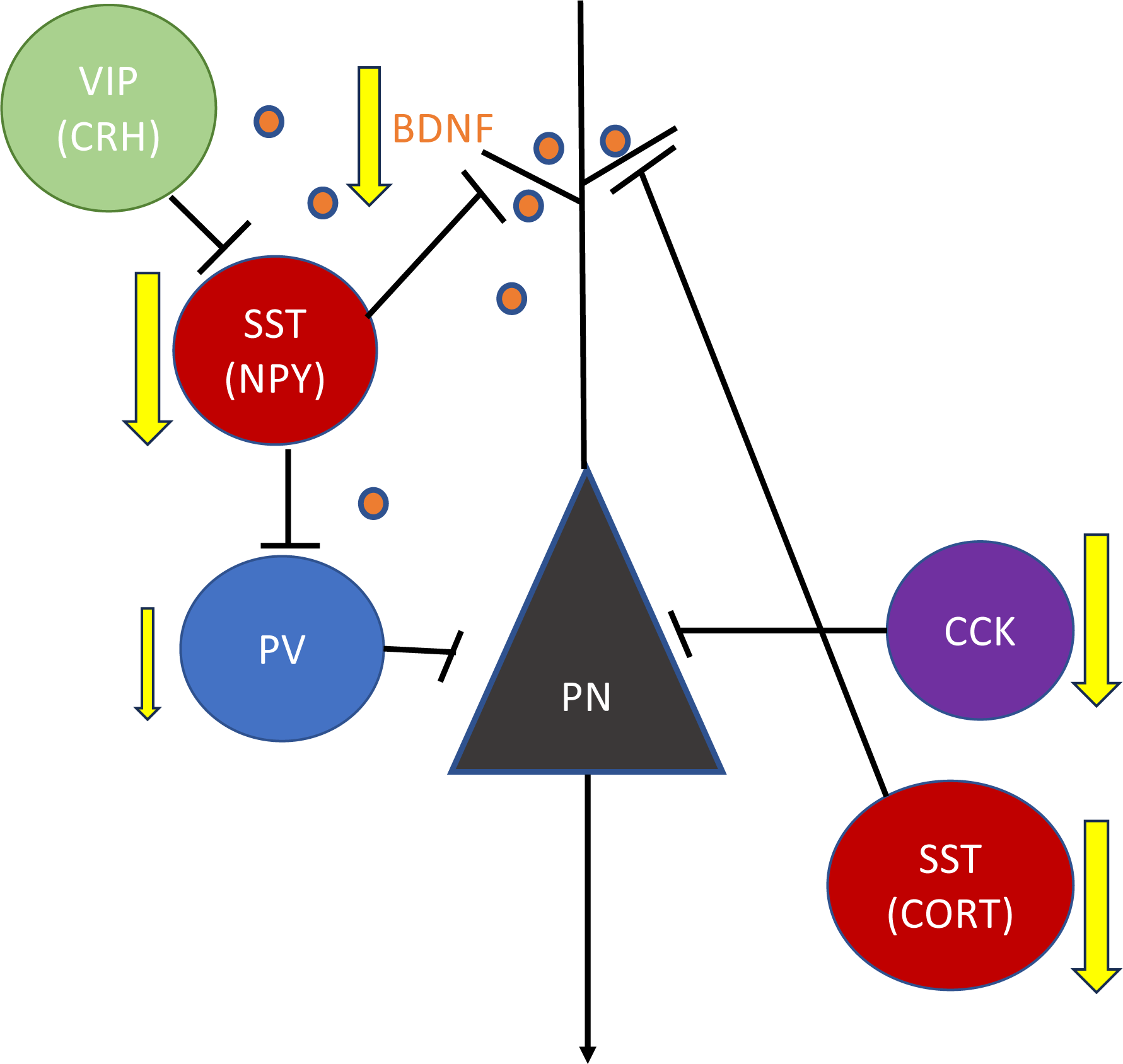
Cortical microcircuit schematic showing the markers affected by LPS-induced inflammation. Pyramidal (PN) neurons are excitatory glutamatergic cells regulated by different inhibitory GABAergic interneurons. Somatostatin (SST) neurons inhibit pyramidal neuron (PN) dendrites. Parvalbumin (PV) neurons inhibit the soma and axon initial segment of pyramidal neurons. Upon stimulus, vasoactive intestinal peptide (VIP) neurons inhibit SST interneuron firing and induce disinhibition of PNs to allow information processing. Cholecystokinin (CCK) neurons mainly innervate the soma of PNs. The neuropeptide corticotropin-releasing hormone (CRH) is mainly co-expressed with VIP interneurons and is important for providing inhibitory inputs to SST neurons. Neuropeptide Y (NPY) and cortistatin (CORT) are commonly co-expressed with SST interneurons. Brain-derived neurotrophic factor (BDNF) is a signaling neuropeptide mainly secreted from PNs and is important for dendritic compartments and GABAergic interneurons (adapted from Fee et al., 2017). Lines with black arrows indicate excitatory input. Lines with black endlines indicate inhibitory inputs. Lines with yellow arrows indicate that the marker decreased with LPS.

### 4.2 BDNF as the putative upstream regulator of reduced GABAergic interneuron markers

Next, we investigated whether LPS-induced inflammation reduces the selective GABAergic interneuron markers downstream from BDNF and found a robust decrease in *Bdnf* expression in the PFC and HPC (Figure 6). Decreased *Bdnf* expression is found in the brain of rodents injected with i.p. IL1-beta or LPS (Guan and Fang, 2006; Lapchak et al., 1993; Schnydrig et al., 2007; Zhang et al., 2016). GABAergic interneurons do not synthesize BDNF and rely on the supply of other cells, specifically PNs (Ernfors et al., 1990; Kohara et al., 2007; Marty et al., 1997). Therefore, lower BDNF secretion is more likely to affect the cell integrity of GABAergic interneurons that express the BDNF receptor tropomyosin receptor kinase B (TrkB), that promotes neuronal plasticity and survival (Sanchez-Huertas and Rico, 2011). The results from all cohorts in both brain regions show that *Bdnf* expression has the highest positive Pearson correlation with *Sst* (r=0.41), *Cort* (r=0.48), and *Npy* (r=0.46) (Figure 6). We recognize a potential for a two-group effect driving a positive correlation. However, the correlation graphs (Supplement figure 1) show substantial group overlap. The positive correlations between *Bdnf* vs. *Sst, Cort*, and *Npy* are consistent with prior findings that BDNF is present on the dendrites of PNs and is known as the upstream regulator of *Sst, Cort,* and *Npy* expression in GABAergic interneurons (Guilloux et al., 2012; Sakata et al., 2009; Tripp et al., 2012). These results suggest that LPS-induced reduction in *Bdnf* may be a critical mediator of the reduced SST interneuron markers. A lower positive correlation of *Bdnf* vs. *Pv* (r=0.33) and *Cck* (r=0.35) may reflect the degree to which additional factors contribute or compensate for the LPS-BDNF effect. In the case of *Pv,* it is shown that this gene is reduced when TrkB function decreases and does not change when *Bdnf* is knocked out (Glorioso et al., 2006; Hashimoto et al., 2005). In contrast, the absence of changes in *Crh* and *Vip* levels and their lack of positive correlation with *Bdnf* can reflect the low-to-no effect of *Bdnf* on those genes, as previously shown that blocking *Bdnf/TrkB* signaling does not reduce *Crh* expression (Oh et al., 2022).

The high levels of IL1-beta and IL-6 in the PFC and HPC (Figure 2) may mediate the reduced *Bdnf* levels by activating the hypothalamic-pituitary-adrenal axis, as previous studies show that glucocorticoids decrease BDNF (Gubba et al., 2004; Hansson et al., 2003). Another possible mechanism for BDNF downregulation may be due to proteinopathy. BDNF is an activity-dependent neuropeptide (Marty et al., 1997), and precursor BDNF undergoes post-translational modifications in the endoplasmic reticulum (ER). In response to LPS, neurons may initially increase their demand on the synthesis of precursor BDNF, exacerbating the ER stress that downregulates mature BDNF synthesis (Tomoda et al., 2021). We measured the mature form of *Bdnf*, but future studies should explore the effect of LPS-induced inflammation on the various BDNF transcripts.

### 4.3 Limitations

LPS-induced inflammation is a well-characterized acute infective model that induces a non-specific systemic inflammation that differs from the unresolved sterile systemic chronic inflammation often associated with MDD (Miller and Raison, 2016). Therefore, LPS-induced inflammation may not reflect the inflammatory changes observed in MDD, which follows a more gradient trajectory. This study focuses on GABAergic interneuron marker expression and correlation downstream of the hallmarks of inflammation following peripheral TLR4-dependent activation and the immune response propagated into the brain. Therefore, further analyses would be required to demonstrate pure causality. This study does not show the functional effect of the reduced GABAergic interneuron markers.

### 4.4 Conclusion

This is the first study investigating the effect of LPS-induced inflammation on markers of the main GABAergic interneurons. It extends our understanding of the interface between the immune and nervous systems. Specifically, the results suggest that inflammation is not a bystander but a putative contributor to the GABAergic interneuron deficits associated with MDD and other neuropsychiatric disorders. This knowledge can help us understand the pathophysiology of MDD with relevance to the role of inflammation on altered GABAergic function to develop potential treatments. Future studies investigating cell-dependent factors may contribute to our understanding of the selective vulnerability of interneurons to inflammation.

## Acknowledgement

We would like to thank CAMH animal facility personnel, including Kristen Fournier and Katrina Deverell, for animal care. We thank Michael Marcotte and Dr. Yashika Bansal for helping with animal brain dissections. We thank Dr. Mouna Banasr for suggesting the cohort 2 experiment design and Mehrab Ali for helping with the manuscript review and submission.

