## Supplement for "LPS-Induced Inflammation Reduces GABAergic Interneuron markers and Brain-derived Neurotrophic Factor in Mouse Prefrontal Cortex and Hippocampus"

### Supplementary Figures List:

**Supplementary Figure 1.** Correlation graphs between *Bdnf* gene and GABAergic interneuron markers (*Sst, Pv, Vip, Cck, Cort, Npy*)

**Supplement figure 1:** Correlation graphs between *Bdnf* gene and GABAergic interneuron markers (*Sst, Pv, Vip, Cck, Cort, Npy*)

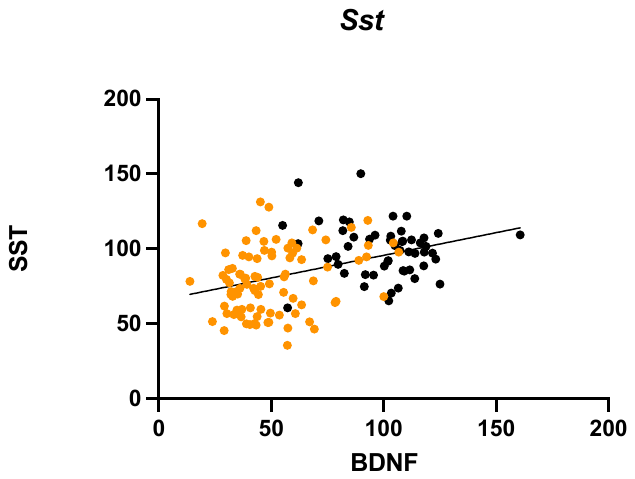

**A**

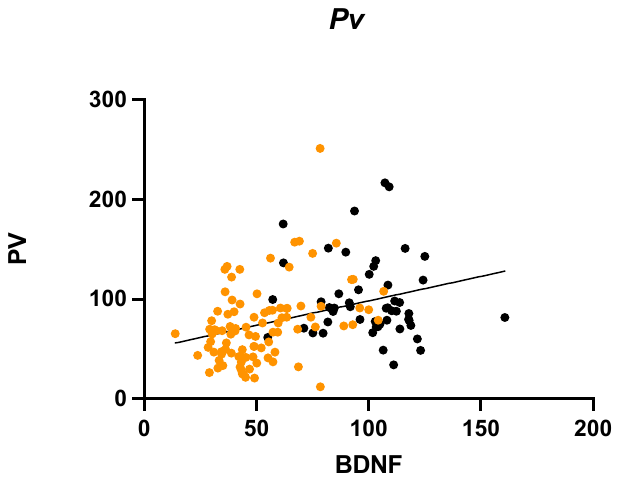

**B**

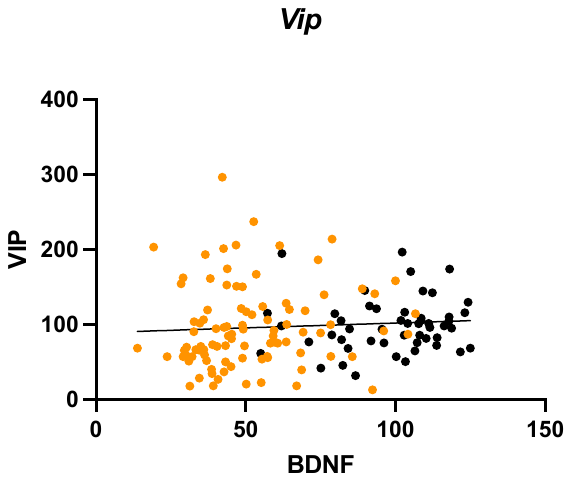

**C**

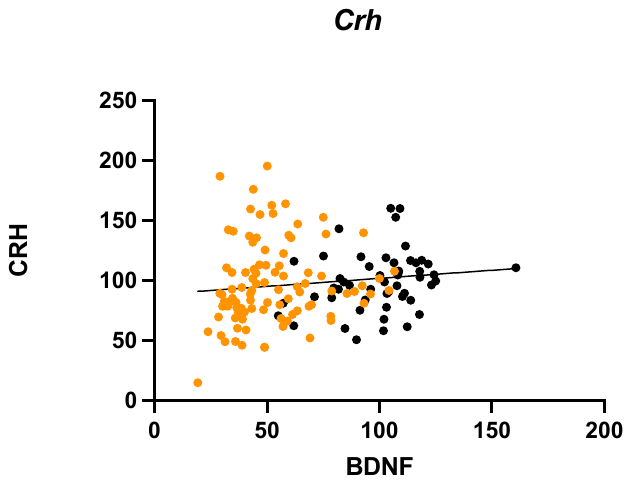

**D**

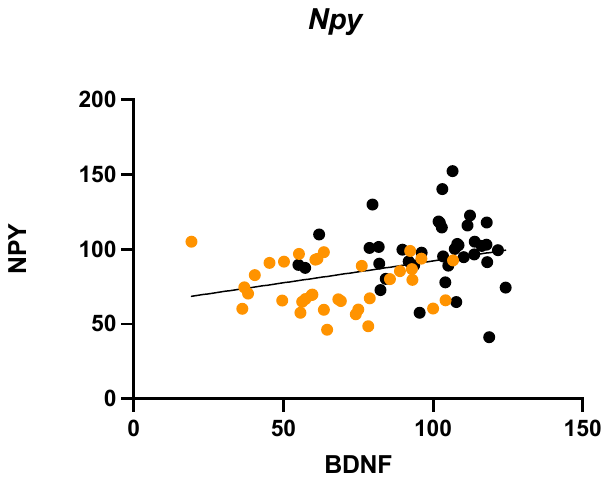

**E**

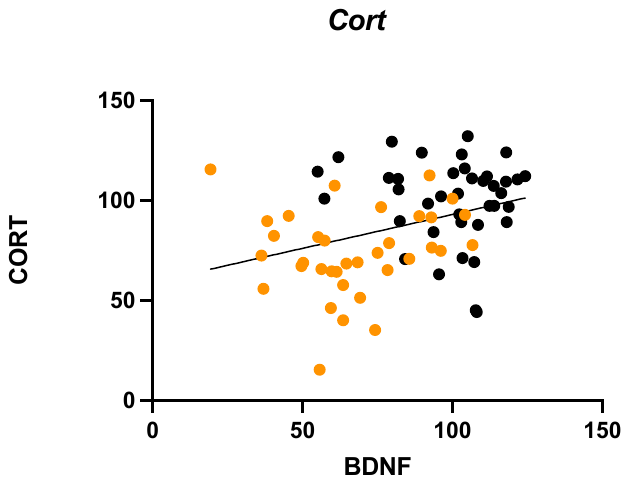

**F**

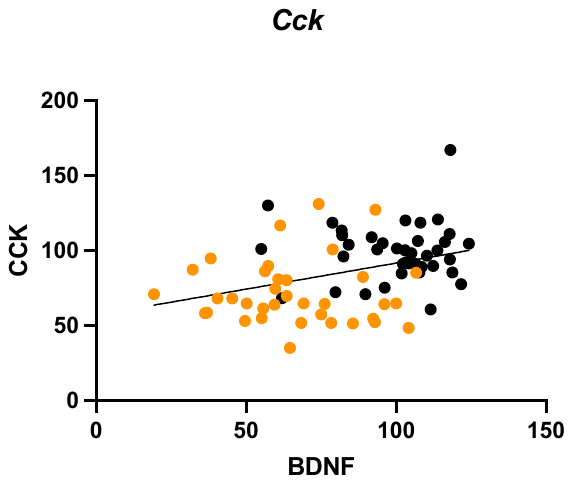

**G**

**SF1: Correlation graphs between *Bdnf* gene and *Sst, Pv, Vip, Crh Npy, Cort,* and *Cck***. **A** *Bdnf* and *Sst* (r=0.41; p<0.0001). **B** *Bdnf* and *Pv* (r=0.33;p<0.0001). **C** *Bdnf* and *Vip* (r=0.08;p=0.31)*.* **D** *Bdnf* and *Crh* (r=0.13;p=0.11). **E** *Bdnf* and *Npy* (r=0.46; p<0.0001). **F** *Bdnf* and *Cort* (r=0.48; p<0.0001). **G** *Bdnf* and *Cck* (r=0.35; p=0.0018). Results are expressed as individual animals. Control mice shown as black circles and LPS mice shown as orange circles.
